## Supplementary File 1 for "Constitutively active receptor ADGRA3 signaling induces adipose thermogenesis"

| **Gene** | **Forward Primer** | **Reverse Primer** |
| --- | --- | --- |
| *Ppargc1α* | TATGGAGTGACATAGAGTGTGCT | CCACTTCAATCCACCCAGAAAG |
| *Pparα* | AGGCTGTAAGGGCTTCTTTCG | GGCATTTGTTCCGGTTCTTC |
| *Ppargγ* | TGTGGGGATAAAGCATCAGGC | CCGGCAGTTAAGATCACACCTAT |
| *Adipoq* | TGTTCCTCTTAATCCTGCCCA | CCAACCTGCACAAGTTCCCTT |
| *Ucp1* | AGGCTTCCAGTACCATTAGGT | CTGAGTGAGGCAAAGCTGATTT |
| *Cidea* | TGCTCTTCTGTATCGCCCAGT | GCCGTGTTAAGGAATCTGCTG |
| *Cox8b* | TGTGGGGATCTCAGCCATAGT | AGTGGGCTAAGACCCATCCTG |
| *Adgra3* | CACGGTCACCCTGATTTTAAGC | GCACCTGGGGCTATCCTACTAA |
| *Gnas* | TGCCTCGGCAACAGTAAGAC | GCCGCCCTCTCCGTTAAAC |
| *Atgl* | GGAGGAATGGCCTACTGAACC | ATCCTCTTCCTGGGGGACAA |
| *Hsl* | CCAGCCTGAGGGCTTACTG | CTCCATTGACTGTGACATCTCG |
| *Actb* | TGTCCACCTTCCAGCAGATGT | AGCTCAGTAACAGTCCGCCTAGA |
| *Hexo* | GCCAGCCTCTCCTGATTTTAGTGT | GGGAACACAAAAGACCTCTTCTGG |
| *mito-16s* | CCGCAAGGGAAAGATGAAAGAC | TCGTTTGGTTTCGGGGTTTC |
| *ACTB* | CATGTACGTTGCTATCCAGGC | CTCCTTAATGTCACGCACGAT |
| *ADGRA3* | GCGTCATTACGGTCTTTGGAA | ACGGCAATTCAAGCGGAGG |
| *UCP1* | AGGTCCAAGGTGAATGCCC | TTACCACAGCGGTGATTGTTC |

**Supplementary File 1. Primer sequences used for qPCR.**
