## Supplementary File 2 for "Constitutively active receptor ADGRA3 signaling induces adipose thermogenesis"

| **Gene ID** | **Gene Name** | **GO Annotation** |
| --- | --- | --- |
| 14748 | ***Gpr3*** | GO0004930: G-protein coupled receptor activity |
| 70693 | ***Adgra3*** | GO0004930: G-protein coupled receptor activity |
| 224792 | ***Adgrf5*** | GO0004930: G-protein coupled receptor activity |
| 170757 | ***Adgrl4*** | GO0004930: G-protein coupled receptor activity |
| 11549 | ***Adra1a*** | GO0004930: G-protein coupled receptor activity |
| 11554 | ***Adrb1*** | GO0004930: G-protein coupled receptor activity |
| 12778 | ***Ackr3*** | GO0004930: G-protein coupled receptor activity |
| 12922 | ***Crhr2*** | GO0004930: G-protein coupled receptor activity |
| 13618 | ***Ednrb*** | GO0004930: G-protein coupled receptor activity |
| 14368 | ***Fzd6*** | GO0004930: G-protein coupled receptor activity |
| 14427 | ***Galr1*** | GO0004930: G-protein coupled receptor activity |
| 242425 | ***Gabbr2*** | GO0004930: G-protein coupled receptor activity |
| 329252 | ***Lgr6*** | GO0004930: G-protein coupled receptor activity |
| 78134 | ***Lpar4*** | GO0004930: G-protein coupled receptor activity |
| 258218 | ***Olfr102*** | GO0004930: G-protein coupled receptor activity |
| 258027 | ***Olfr1385*** | GO0004930: G-protein coupled receptor activity |
| 258334 | ***Olfr1396*** | GO0004930: G-protein coupled receptor activity |
| 634104 | ***Olfr287*** | GO0004930: G-protein coupled receptor activity |
| 259010 | ***Olfr393*** | GO0004930: G-protein coupled receptor activity |
| 258816 | ***Olfr466*** | GO0004930: G-protein coupled receptor activity |
| 259097 | ***Olfr558*** | GO0004930: G-protein coupled receptor activity |
| 259062 | ***Olfr667*** | GO0004930: G-protein coupled receptor activity |
| 258469 | ***Olfr90*** | GO0004930: G-protein coupled receptor activity |
| 258825 | ***Olfr975*** | GO0004930: G-protein coupled receptor activity |
| 18441 | ***P2ry1*** | GO0004930: G-protein coupled receptor activity |
| 94226 | ***S1pr5*** | GO0004930: G-protein coupled receptor activity |
| 21337 | ***Tacr2*** | GO0004930: G-protein coupled receptor activity |

**Supplementary File 2. Gene annotation for screened genes.** The 1134 screened genes were annotated by David database and 27 of these genes were identified as G-protein coupled receptor encoding gene.
